# Widespread variation in *EDNRB2* is associated with diverse melanin loss phenotypes across avian species

**DOI:** 10.1101/2025.06.13.659395

**Authors:** Emily T. Maclary, Michael D. Shapiro

**Author notes:** Author for Correspondence Emily T. Maclary, School of Biological Sciences, 257 South 1400 East, Salt Lake City, UT 84112 USA.

## Abstract

Plumage pigmentation plays critical roles in survival and reproductive success in birds, from providing camouflage and thermoregulation to mediating elaborate mating displays. The genetic and developmental origins of diverse plumage pigmentation patterns remain incompletely understood in part due to limited intraspecific variation and high levels of genetic divergence between distantly related species. Domestic avian species are more tractable models for understanding the genetic architecture of plumage pigmentation, but the relevance of domestic phenotypes to plumage patterns observed in the wild is not clear. Here, we used comparative genomic approaches to examine coding variation in *EDNRB2,* a candidate gene associated with loss of plumage melanin in several species, in representative genomes from a diverse array of wild and domestic birds. We found widespread coding variation in *EDNRB2* and in other pigmentation genes with limited pleiotropic roles in development. We also found that *EDNRB2-* mediated melanin loss may play a critical role in establishing bright non-melanin plumage colors. This work highlights *EDNRB2* as a key candidate gene for mediating the development of both interspecific and intraspecific plumage variation and demonstrates the applicability of findings in domestic species to understanding avian plumage patterning more broadly.

## INTRODUCTION

Birds display striking differences in plumage color and pattern across species. In a natural setting, plumage pigmentation can impact diverse processes that influence survival and reproductive success, including thermoregulation, camouflage, mate choice, and communication (Hubbard et al. 2010; Orteu and Jiggins 2020; Price-Waldman and Stoddard 2021). Longstanding interest in understanding the genetic architecture of avian plumage pigmentation and patterning has inspired numerous studies in domestic and wild species. Domestic animals, including birds, are often bred for specific pigment phenotypes, and studies of domestic populations have led to critical insights about the genetic control of pigment production and distribution (Cieslak et al. 2011; Domyan et al. 2014; Cooke et al. 2017; Vickrey et al. 2018; Bruders et al. 2020; Price-Waldman and Stoddard 2021; Bovo et al. 2023)

While studying intraspecific variation in domestic species is a tractable approach to identifying and understanding the genetic architecture of pigment-related traits, some genetic variants affecting pigmentation are associated with deleterious pleiotropic effects such as eye defects, hearing loss, altered growth, and osteoporosis, which would be maladaptive in wild species (Kawaguchi et al. 2001; Hoekstra 2006; Minvielle et al. 2010; Cieslak et al. 2011; Bruders et al. 2020; Varga et al. 2020). Pigmentation variants are maintained in domestic animals despite these effects due to selection by breeders for novel color phenotypes (Cieslak et al. 2011), raising questions about the relevance of candidate genes in domestic species for understanding pigmentation diversity in the wild. We can hypothesize that the mechanisms that give rise to pigmentation diversity among wild species are similar; however, relatively few studies have examined the genetics of coloration or patterning in wild species, and pleiotropic effects may limit whether some of the genes implicated in domestic species can realistically underlie variation in the wild.

Though wild birds show spectacular diversity in plumage patterning across species, they often lack substantial intraspecific diversity in pigmentation patterning, thereby making them poor models to discover the genetic basis of variation. Comparing genomes between species with pigmentation differences is also challenging due to high levels of genetic divergence, much of which is unlikely to affect plumage color traits. Recently, however, several genomic studies have successfully identified genes associated with plumage variation in wild species by examining hybrid zones, closely related recent radiations, or populations with intraspecific variation among populations. For example, examining a hybrid zone between subspecies of white wagtails (*Motacilla alba*) with divergent head coloration showed that the agouti signaling protein gene *ASIP* is associated with variation in plumage coloration (Semenov et al. 2021), while analysis of two closely related corvid species showed that *NDP* is associated with plumage variation between all-black carrion crows and black and gray hooded crows (Poelstra et al. 2014; Knief et al. 2019). Other recent studies of hybrid zones identified *CYP2J19* as a candidate associated with a shift between red and yellow carotenoid coloration in northern flickers and *Pogoniulus* tinkerbirds (Kirschel et al. 2020; Aguillon et al. 2021). Analysis of a rapid radiation of Capuchino seedeaters identified several genomic regions containing known pigment genes are associated with color variation in a recent radiation, including *ASIP, KITL, SLC45A2, TYRP1,* and *MLANA* (Campagna et al. 2017). Notably, all these aforementioned genes have also been identified as candidate plumage pigmentation genes in domestic species (Domyan et al. 2014; Lopes et al. 2016; Mundy et al. 2016; Vickrey et al. 2018; Yu et al. 2019; Bruders et al. 2020). These shared candidate genes identified in wild and domestic populations indicate that, despite the negative pleiotropies associated with some pigment phenotypes in domestic species, studies of domestic populations are a promising and tractable way to discover general genetic mechanisms that underlie plumage pigmentation diversity.

We recently identified an allelic series at the *EDNRB2* locus in the domestic pigeon (*Columba livia*) associated with a suite of depigmentation phenotypes known as piebalding (Maclary et al. 2021; Maclary et al. 2023). *EDNRB2* has been implicated in plumage pigmentation variation in several other domestic birds, including chicken, duck, goose, quail, peafowl, and zebra finch (Miwa et al. 2007; Kinoshita et al. 2014; Li et al. 2015; Wu et al. 2017; Xi et al. 2020; Xi et al. 2021; Ouyang et al. 2022; Rodríguez-Martínez 2022; Nannan et al. 2023; Wang et al. 2023). Phenotypes associated with *EDNRB2* variants vary widely, and include mottling, patches of white plumage (piebalding), and complete loss of plumage melanin. The types of *EDNRB2* variants associated with melanin loss include coding, noncoding, and structural variants. Plumage patterns with white patches where melanin is absent are also common in myriad wild bird species across multiple orders; however, no studies to date have identified *EDNRB2*-linked plumage variation in wild avian species. Given the repeated involvement of *EDNRB2* in piebalding in genetically tractable domestic species and the similar melanin-loss phenotypes that evolved in many wild birds, we hypothesized that this gene is a viable candidate for controlling interspecific variation in pigmentation patterning in addition to contributing to intraspecific plumage diversity in domestic populations.

The endothelin signaling pathway evolved through multiple rounds of gene duplication, and most vertebrates have two endothelin B receptor genes, *EDNRB* and *EDNRB2*, that are critical for neural crest cell migration during embryogenesis (Pla and Larue 2003; Pla et al. 2005; Harris et al. 2008; Braasch et al. 2009). Different endothelin B receptor genes are expressed in different neural crest cell populations: *EDNRB2* is expressed in migrating melanoblasts and postmigratory melanocytes, where it regulates pigment cell migration and differentiation, while *EDNRB* is expressed in non-pigment neural crest cells, including skeletal and trunk neural crest. Mammals have only one endothelin B receptor gene, *EDNRB*, that controls both pigment and non-pigment cell neural crest migration, and mutations are associated with both melanin loss and enteric nervous system defects such as aganglionic megacolon (Metallinos et al. 1998; Pla and Larue 2003; Reissmann and Ludwig 2013). The subfunctionalization of endothelin receptors in birds makes these genes intriguing candidates for examining sequence diversity and mutation burden across species: *EDNRB* controls conserved developmental processes critical for peripheral nervous system development and mutations are likely to be detrimental, while *EDNRB2* specifically regulates pigment cell migration, differentiation, and survival, which vary widely among species with divergent plumage patterns. Coding changes in *EDNRB2* could result in reduced protein function, including null alleles with total loss of neural crest-derived melanin, or regulatory changes could alter gene expression, but due to the subfunctionalization of *EDNRB2,* no other neural crest-derived cells would be impacted. With these functional differences in mind, we hypothesized that *EDNRB* should be under strong purifying selection among birds, while selective pressure on *EDNRB2* would be comparatively relaxed in some or all clades.

Here, we examine coding sequence variation in endothelin B receptor genes in the representative genome assemblies from a phylogenetically and phenotypically diverse array of domestic and wild avian species. We show that avian orthologs of *EDNRB2* have an increased nucleotide substitution rate compared to *EDNRB,* and have accumulated significantly more synonymous and nonsynonymous mutations over evolutionary time. Through comparisons with other pigmentation genes, we find that this elevated mutation rate is characteristic of genes with limited pleiotropy. Within published reference genome assemblies, we identify several coding variants in *EDNRB2* that are likely to impact protein function, including previously uncharacterized variants in domestic species that are associated with melanin loss phenotypes and variants in wild species that may drive species-specific melanin loss. In addition, we find that regional melanin loss is likely essential for establishing bright patches of non-melanin plumage coloration, including carotenoid and psittacofulvin-based pigments.

## RESULTS

### Coding variation in *EDNRB2* is widespread among avian species

Due to the subfunctionalization of *EDNRB2* and the presence of diverse patterns of melanin loss in both wild and domestic birds, we hypothesized that *EDNRB2* would show higher levels of sequence diversity across avian species compared to *EDNRB*. To examine coding sequence diversity, we aligned the coding portions of *EDNRB* and *EDNRB2* mRNA sequences from 85 avian species representing 59 families, including 11 species with domestic populations (fig. 1A, supplementary table S1). In both *EDNRB* and *EDNRB2* gene models, we found some variability in annotated coding sequence length (supplementary fig. S1). Variation in length is predominantly driven by variation in the region encoding the extracellular N-terminus of the protein (supplementary fig. S2). For consistency and direct comparability across taxa, we standardized coordinates for analysis to the coding sequence of the chicken *EDNRB2* annotation [NM_001001127.2] and its associated protein model [NP_001001127.1].

**Figure 1:**
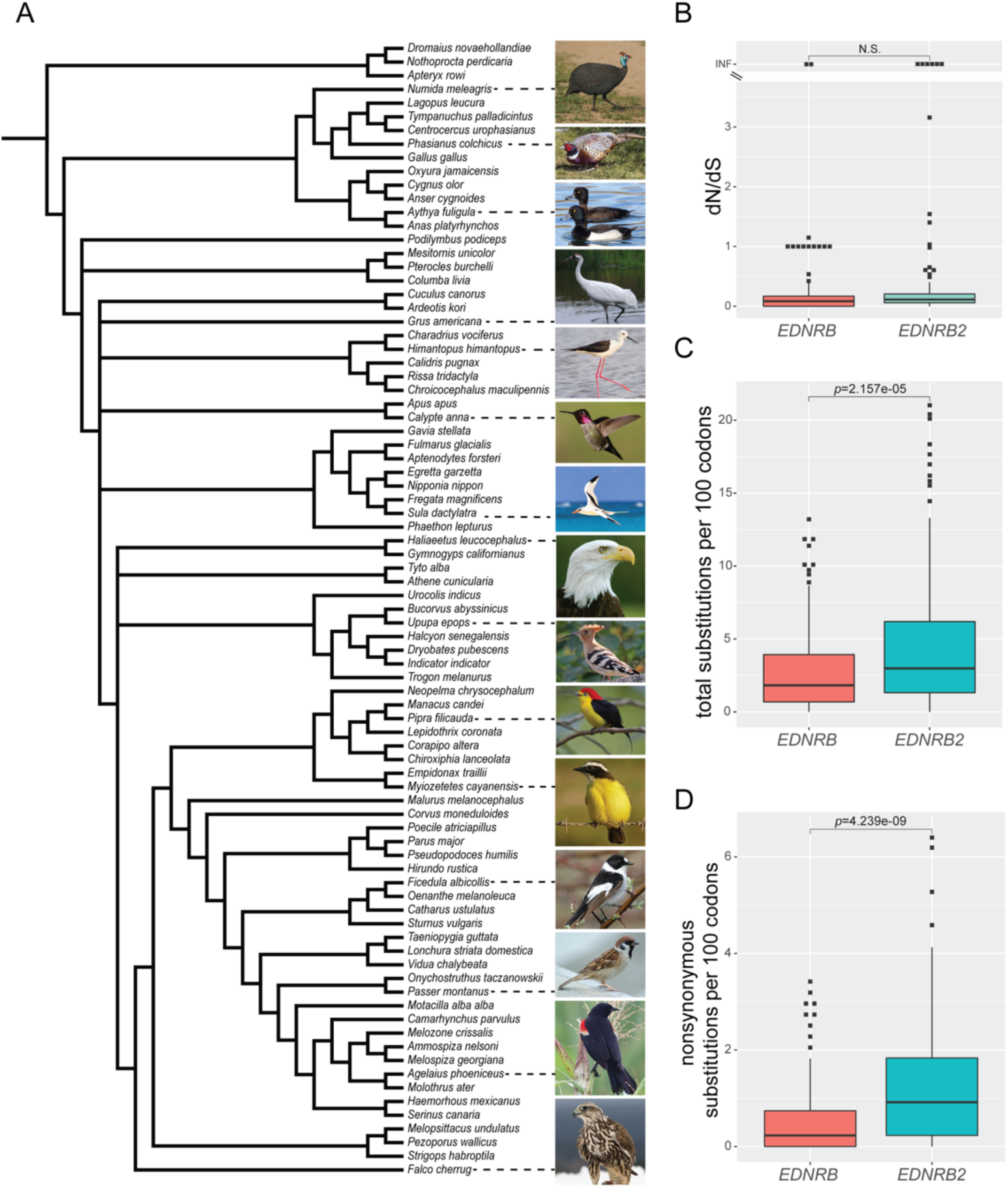
Comparison of total and nonsynonymous substitution rates across 85 species in *EDNRB* and *EDNRB2*. (A) Cladogram showing relationships between the 85 species used for comparative analysis of *EDNRB* and *EDNRB2.* Photographs illustrate diversity in plumage phenotypes from a subset of species, including several species exhibiting plumage patches that lack melanin. Image credits and creative commons license details are in supplementary table S6. (B) Boxplots of per-branch nonsynonymous to synonymous nucleotide substitution ratios (dN/dS). Distribution of per-branch dN/dS does not significantly differ between *EDNRB* and *EDNRB2*. Boxes span from the first to third quartile of each data set, with lines indicating the median. Whiskers span up to 1.5x the interquartile range. (C) Boxplot showing the distribution of total substitutions per 100 codons for each cladogram branch. Total substitutions in *EDNRB2* are significantly higher than in *EDNRB*. (D) Boxplot showing the distribution of nonsynonymous substitutions per 100 codons for each cladogram branch. Nonsynonymous substitutions in *EDNRB2* are significantly higher than in *EDNRB*.

We quantified nonsynonymous and synonymous substitutions on each branch of a species tree inferred from published phylogenies (fig. 1A) (Sun et al. 2017; Oliveros et al. 2019; Kuhl et al. 2020; Braun and Kimball 2021; Chen et al. 2021; Smith et al. 2022). We found that the ratio of nonsynonymous to synonymous substitutions (dN/dS) per branch did not differ significantly between *EDNRB* and *EDNRB2*, and most branches have a dN/dS < 1, indicating some level of purifying selection (fig. 1B, supplementary table S2). A small number of branches showed infinite dN/dS values due to the presence of a single nonsynonymous substitution and no synonymous substitutions. While the ratio of nonsynonymous to synonymous substitutions did not differ significantly, we found that *EDNRB2* had a significantly elevated number of total substitutions and nonsynonymous substitutions (fig. 1C-D).

We next examined the locations of nonsynonymous substitutions within *EDNRB* and *EDNRB2*. We found that sequence variants in both genes were concentrated in the regions that encode the extracellular N-terminal domain, with 83% of nonsynonymous substitutions in *EDNRB* and 72% of nonsynonymous substitutions in *EDNRB2* occurring in this region (fig. 2). For *EDNRB*, all identified indels within the coding sequence mapped to the extracellular N-terminus, and all except two coding indels in *EDNRB2* mapped to the extracellular N-terminus. Despite its importance for endothelin B receptor localization and function, we found that the regions of *EDNRB* and *EDNRB2* that encode the extracellular N-terminal domain are highly variable in birds, both in length and sequence content. Most species included in these alignments have at least regional plumage melanin, indicating some level of EDNRB2 functionality. As a result, it is challenging to determine the possible impacts of N-terminal variation from alignment of divergent genomes, though we conclude that variation in the extracellular N-terminus is generally tolerated in both proteins.

**Figure 2:**
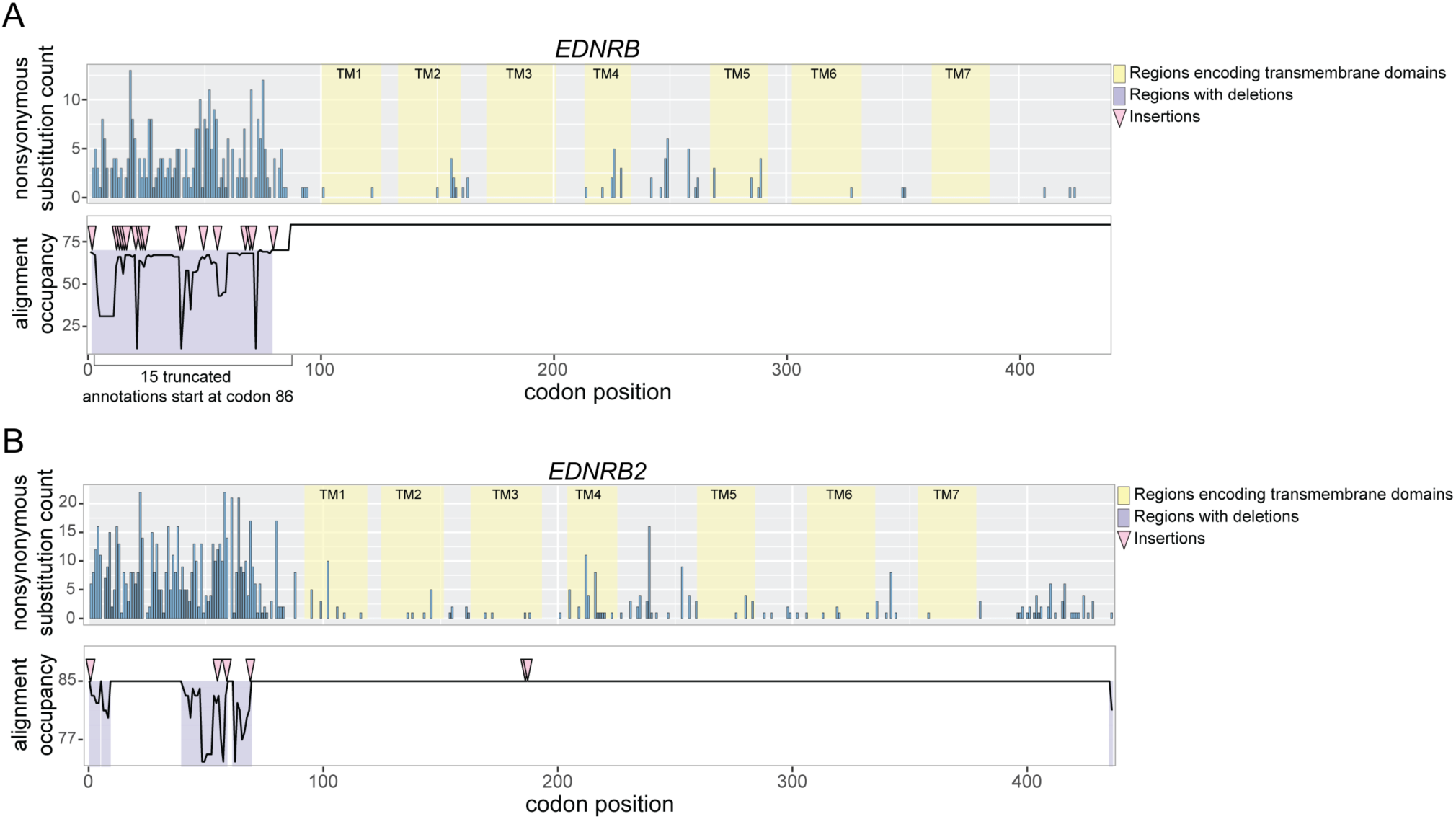
Distribution of nonsynonymous substitutions and indels in *EDNRB* and *EDNRB2*. Plots illustrating the distribution of nonsynonymous substitutions in reference annotations for *EDNRB* (A) and *EDNRB2* (B). Codon positions are based on chicken as a reference sequence. In upper panels, blue bars show the count of nonsynonymous substitution events per codon. Yellow highlights show regions encoding transmembrane domains. In lower panels, a line plot shows alignment occupancy, or the number of species that have any aligned sequence spanning that codon. Lower occupancy indicates the presence of a deletion at that site in one or more species. Pink triangles mark sites where one or more species has an insertion relative to the chicken reference sequence. Several *EDNRB* annotations are truncated relative to chicken, missing the first 85 codons. This region that is absent from 15/85 annotations is marked with a bracket.

Beyond the N-terminal extracellular domain, the difference in amino acid substitution rates is also striking. In *EDNRB*, only 17% of nonsynonymous substitutions are outside of the region encoding the N-terminal extracellular domain, but in *EDNRB2*, 28% of nonsynonymous substitutions are outside of the region encoding this domain. In sum, we find that *EDNRB2* shows an increased nonsynonymous substitution rate compared to *EDNRB*, consistent with our hypothesis, and we see stark differences in the distribution of substitutions across the regions encoding different protein domains.

### Synonymous and nonsynonymous substitutions rates vary across pigmentation genes

We next wanted to evaluate coding mutation burden in pigmentation genes more broadly since, like *EDNRB2*, several other genes have been implicated repeatedly in plumage or coat color polymorphisms in domestic species. We hypothesized that pleiotropy may limit coding sequence diversity in some pigmentation genes but, like *EDNRB2*, other genes with pigmentation-specific functions may show high levels of interspecific sequence variation. We assessed variation in coding sequences for *EDNRB2* and five other genes previously linked to pigmentation variation in numerous domestic species: *ASIP, MC1R, MITF, SOX10,* and *TYRP1.* Here, we chose a subset of 35 avian species representing 35 families for analysis (supplementary table S1). We aligned the coding portion of annotated mRNA sequences and quantified synonymous and nonsynonymous mutations for each branch of the phylogeny. We found that the distribution of dN/dS values is significantly higher in *ASIP* than in other genes, suggesting that *ASIP* may be evolving rapidly under diversifying selection in more species than the other pigmentation genes we assessed (fig. 3A, supplementary table S3).

**Figure 3:**
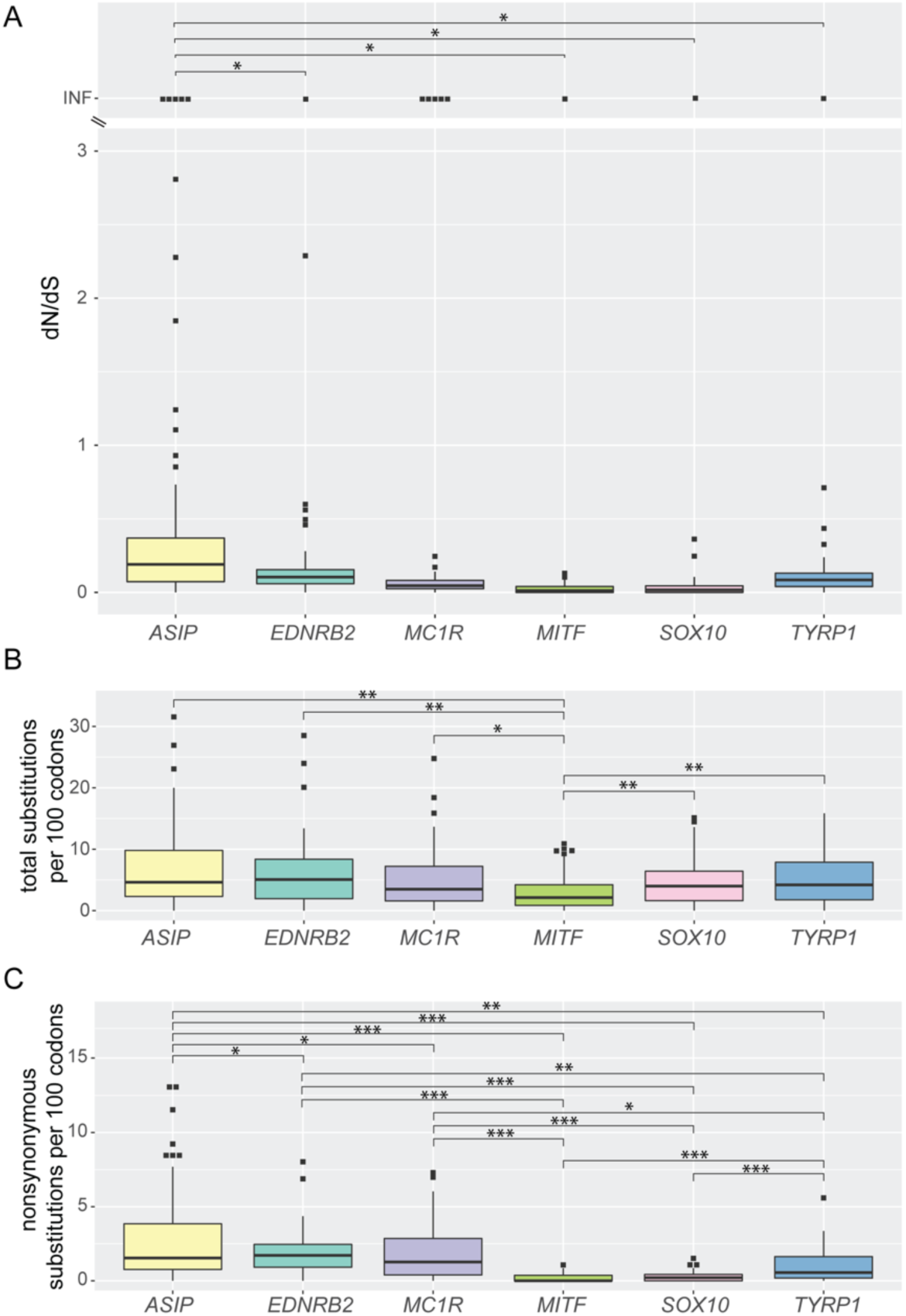
Comparison of substitution rates across six pigment-associated genes. Boxplots illustrating (A) dN/dS ratio, (B) total number of substitutions per 100 codons, and (C) nonsynonymous substitutions per 100 codons for six pigment-associated genes. Substitutions were calculated from alignments of 35 species representing a subset of the species used for alignments in fig. 1. Boxes span from the first to third quartile of each data set, with lines indicating the median. Whiskers span up to 1.5x the interquartile range. *,0.05> *p ≥*0.005; **,0.005> *p ≥*5e-5; ***, *p* <5e-5, two-tailed t-test.

While dN/dS values did not significantly differ among other pigmentation genes, we again found significant variation in the total number of substitutions and number of nonsynonymous substitutions (fig. 3B-C). The total number of substitutions per 100 codons was significantly lower in *MITF* compared to other genes, and the numbers of nonsynonymous substitutions per 100 codons were significantly lower for both *MITF* and *SOX10*. Notably, mutations affecting both *MITF* and *SOX10* – both of which encode transcription factors with multiple roles in development – are associated with well-documented negative pleiotropic effects in domestic birds and other vertebrates, including microphthalmia, blindness, osteoclast defects, hearing loss, reproductive changes, aganglionic megacolon, and lethality (Kawaguchi et al. 2001; Stritzel et al. 2009; Pingault et al. 2010; Reissmann and Ludwig 2013). In contrast, mutations in *ASIP, EDNRB2, MC1R,* and *TYRP1* are common in domestic and wild vertebrates and are primarily linked to pigmentation traits. Nevertheless, a subset of *TYRP1* variants is linked to progressive hearing loss, a subset of *ASIP* alleles are associated with metabolic changes and increased tumorigenesis in mice, and some *MC1R* variants are linked to melanoma risk, immune response, and analgesia (Johnson and Jackson 1992; Voisey and Daal 2002; Mogil et al. 2005; Fierro-Castro et al. 2022), indicating that viable mutations with pleiotropic effects occur, though they may not be common.

For the six pigmentation genes we evaluated, we found that coding variation between species is higher in genes where mutations typically have fewer obvious pleiotropic effects, including *EDNRB2.* We cannot rule out that mutations may occur at similar rates in the other pigmentation genes, but pleiotropic effects ranging from lethality to reduced reproductive success would be selected against, thereby eliminating our changes of recovering them in genome assemblies. However, we also see a shift in the number of synonymous substitutions per 100 codons for genes with limited pleiotropy. The high number of nonsynonymous substitutions per 100 codons identified in *EDNRB2* is comparable to that of *MC1R,* and coding changes in *MC1R* are associated with plumage diversity in several wild avian species (Theron et al. 2001; Mundy et al. 2004; Mundy 2005; Corso et al. 2016; Campagna et al. 2022; Ono et al. 2024). We conclude that pleiotropy likely constrains the genetic architecture of plumage pigment variation, and that limited pleiotropy and high rates of coding variation in *EDNRB2* make it a prime candidate for contributing to plumage pigment variation both within and between avian species.

### Avian genomes harbor *EDNRB2* substitutions predicted to impact protein function

We identified numerous nonsynonymous mutations in *EDNRB2* in our 85-species analysis. However, it is unclear if any of these mutations are linked to changes in plumage pigmentation. To screen for substitutions that may impact protein function, we first aligned predicted protein sequences for all three endothelin receptors – EDNRA, EDNRB, and EDNRB2 – from a diverse set of tetrapods (human, mouse, chicken, western clawed frog, common wall lizard, and green sea turtle) and identified fully conserved amino acid residues. We also used the NCBI Conserved Domain Database to identify transmembrane domains and putative ligand binding sites. We then examined the distribution of amino acid substitutions and indels throughout the EDNRB2 protein and compared the sites of mutations to predicted functional domains and evolutionarily conserved residues. While 72% (288 out of 401) of amino acid substitutions are in the highly variable N-terminal region, we also identified 53 amino acid substitutions within transmembrane domains and 6 at positions predicted to be involved in ligand binding. Eight of the variant sites were 100% conserved in our tetrapod endothelin receptor alignment. Of these 8 conserved residues, 5 are in transmembrane domains, one is in an extracellular domain, and 2 are in intracellular domains (table 1). Like the amino acid substitutions, most indels were also within the extracellular N-terminal region, except for two in-frame insertions of single amino acids in a stretch of well-conserved sequence that spans the intracellular domain between TM3 and TM4 and the start of the fourth transmembrane domain.

**Table 1:**
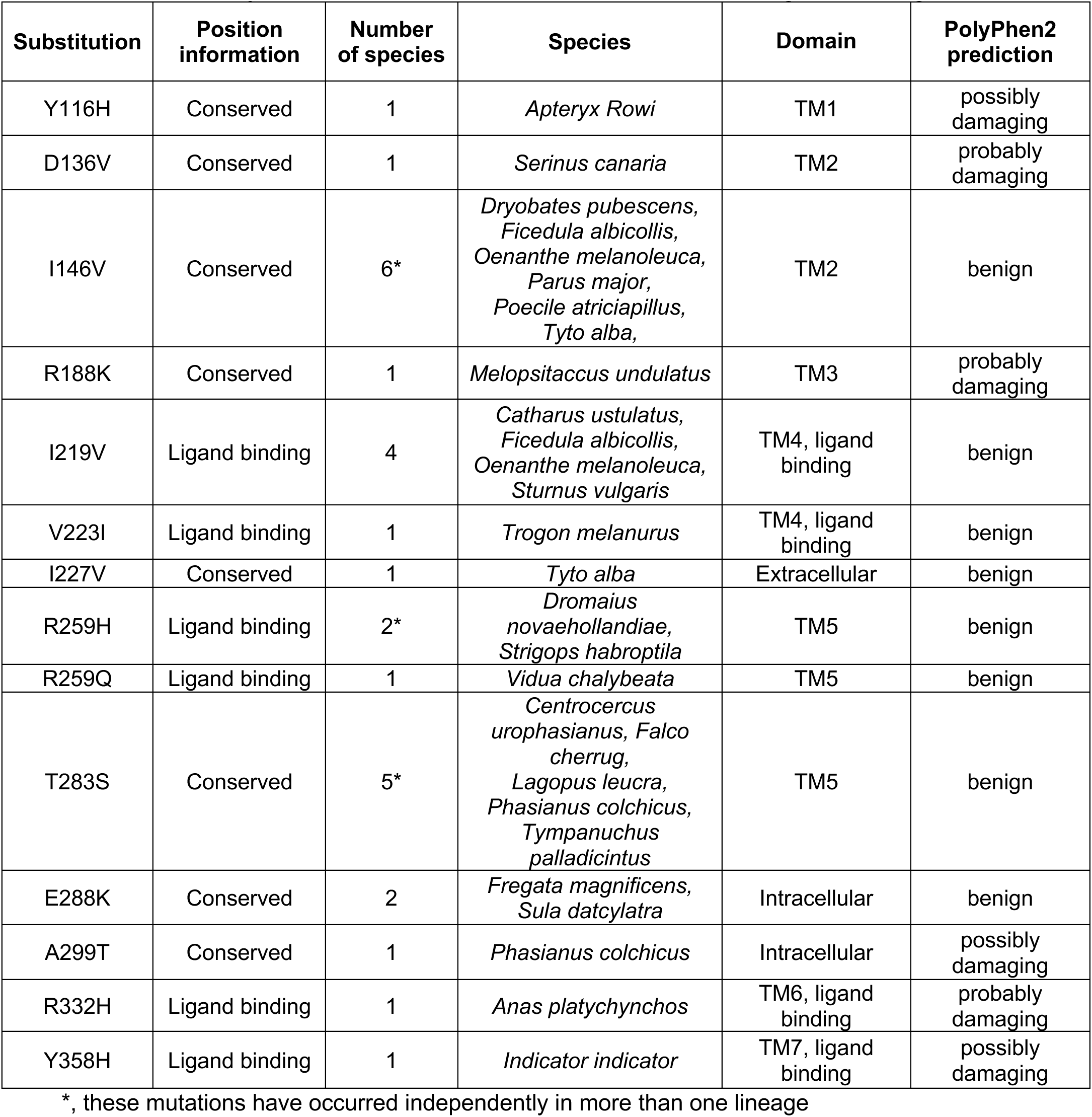
Summary of amino acid substitutions at conserved and ligand binding residues.

| Substitution | Position information | Number of species | Species | Domain | PolyPhen2 prediction |
| --- | --- | --- | --- | --- | --- |
| Y116H | Conserved | 1 | *Apteryx Rowi* | TM1 | possibly damaging |
| D136V | Conserved | 1 | *Serinus canaria* | TM2 | probably damaging |
| I146V | Conserved | 6* | *Dryobates pubescens*,<br>*Ficedula albicollis*,<br>*Oenanthe melanoleuca*,<br>*Parus major*,<br>*Poecile atriciapillus*,<br>*Tyto alba*, | TM2 | benign |
| R188K | Conserved | 1 | *Melopsitaccus undulatus* | TM3 | probably damaging |
| I219V | Ligand binding | 4 | *Catharus ustulatus*,<br>*Ficedula albicollis*,<br>*Oenanthe melanoleuca*,<br>*Sturnus vulgaris* | TM4, ligand binding | benign |
| V223I | Ligand binding | 1 | *Trogon melanurus* | TM4, ligand binding | benign |
| I227V | Conserved | 1 | *Tyto alba* | Extracellular | benign |
| R259H | Ligand binding | 2* | *Dromaius novaehollandiae*,<br>*Strigops habroptila* | TM5 | benign |
| R259Q | Ligand binding | 1 | *Vidua chalybeata* | TM5 | benign |
| T283S | Conserved | 5* | *Centrocercus urophasianus*, *Falco cherrug*,<br>*Lagopus leucra*,<br>*Phasianus colchicus*,<br>*Tympanuchus palladicintus* | TM5 | benign |
| E288K | Conserved | 2 | *Fregata magnificens*,<br>*Sula dactylatra* | Intracellular | benign |
| A299T | Conserved | 1 | *Phasianus colchicus* | Intracellular | possibly damaging |
| R332H | Ligand binding | 1 | *Anas platychynchos* | TM6, ligand binding | probably damaging |
| Y358H | Ligand binding | 1 | *Indicator indicator* | TM7, ligand binding | possibly damaging |

279 \*, these mutations have occurred independently in more than one lineage

To prioritize variants for follow up, we opted to first examine the 14 amino acid substitutions and 2 insertions that affect either conserved or ligand-binding residues (table 1). Five of the substitutions were present in multiple bird species, with three occurring independently in different lineages (table 1). Nine amino acid substitutions and both insertions were each unique to a single species. We evaluated these substitutions using PolyPhen2 (Adzhubei et al. 2013) and found that eight variants, including all the variants present in multiple species, were predicted to be benign. Six variants were predicted to be “possibly damaging” or “probably damaging” (table 1). These six putative damaging substitutions and the two insertions in the intracellular domain between TM3 and TM4 are discussed below.

### A novel *EDNRB2* coding variant is associated with loss of melanin in canaries

Wild common canaries *(Serinus canaria)* are yellow-green with brown markings throughout the face, body, and wings (fig. 4A), and domestic populations have been bred for centuries and are common as pets. Domestic canaries have many color morphs, including several that exhibit melanin loss, like the Lipochrome yellow form (fig. 4B). In canaries, melanin pigments are required for the brown and green coloration seen in wild-type plumage. Yellow plumage coloration is derived from dietary carotenoids, and loss of carotenoids in domestic lipochrome canaries – which also lack plumage melanins – produces white plumage (Toomey et al. 2017).

**Figure 4:**
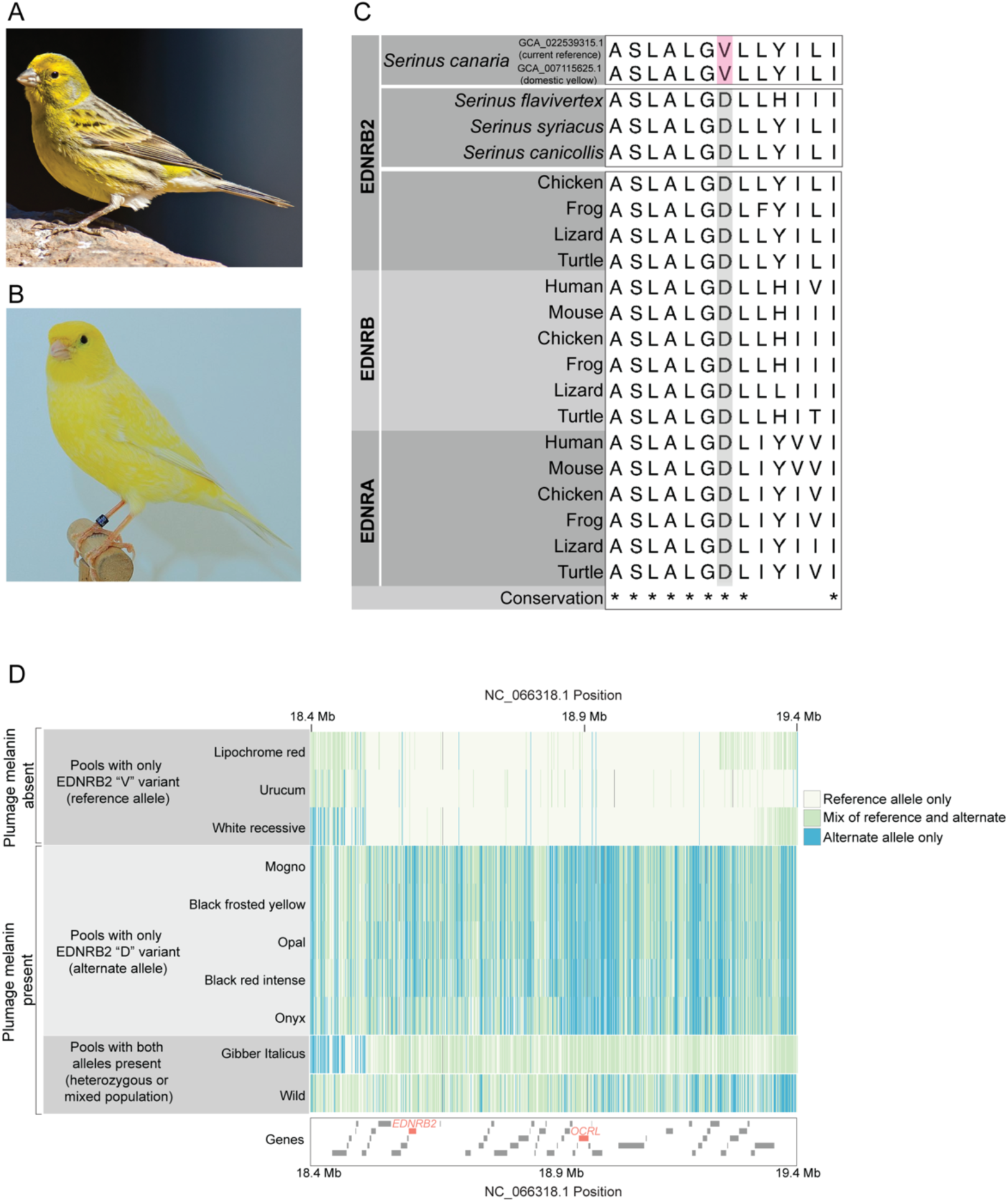
Identification of a substitution in the *Serinus canaria* reference genome associated with melanin loss. Representative images of a wild-type Atlantic canary (A) and a domestic Lipochrome yellow morph that lacks melanin (B). Image credits and creative commons license details are in supplementary table S6. (C) Regional alignments highlighting the D136V mutation in *S. canaria* genomes (pink) compared to the reference genomes of additional *Serinus* species and a multi-species alignment of EDNRA, EDNRB, and EDNRB2 from diverse vertebrates. * indicates amino acids that are 100% conserved in the vertebrate endothelin receptor alignment. (D) Plot of genotypes surrounding the *EDNRB2* locus in pooled sequencing from *S. canaria* samples. Each row shows a sequencing pool from individuals with a shared phenotype, and each vertical line a SNP position colored by genotype relative to the reference genome. Approximate locations of genes are illustrated below, with *EDNRB2* and *OCRL* highlighted in red. For each pool, relevant phenotype information and genotypes for the candidate SNP that causes the D136V substitution are noted on the left-hand side.

We identified a nonsynonymous substitution leading to an amino acid change at a conserved amino acid in transmembrane domain 2 (D136V) in a canary reference genome [GCA_022539315.1]. We screened for this amino acid substitution in three additional wild *Serinus* species with published reference genomes and found that none shared the D136V substitution (fig. 4C, supplementary table S4). We next examined the other two available reference genome assemblies from the common canary and found that the D136V substitution was present in one of them [GCA_007115625.1], which is derived from a domestic individual. The presence of the variant could not be determined in the other reference [GCA_000534875.1] due to gaps in the genomic region containing *EDNRB2*. Based on available data, we hypothesized that the D136V mutation is associated with melanin loss in domestic canaries.

To test this possibility, we examined previously published resequencing data from domestic canaries with different plumage colors. Prior work generated pooled sequencing from domestic canary populations to map the genetic architecture of various pigmentation traits (Toomey et al. 2017; Gazda et al. 2020; Bovo et al. 2023). Three of these sequencing pools are from breeds that exhibit total melanin loss: Lipochrome red, which has bright red plumage; Urucum, which is a lipochrome color that additionally shows carotenoid coloration on bare parts like feet; and White recessive, which has white plumage with no carotenoid or melanin pigmentation (Toomey et al. 2017; Gazda et al. 2020; Bovo et al. 2023). We found that all three populations were fixed for the SNP that causes the D136V substitution and shared an approximately 700-kb haplotype at the *EDNRB2* locus (fig. 4D). Pooled sequences from five populations with melanin-containing plumage morphs (Mogno, Black frosted yellow, Opal, Black red intense, and Onyx) did not have the D136V substitution.

We found that both alleles were present in the resequenced “Gibber Italicus” and “Wild” pools of canaries. In the Wild pool, 25% of mapped reads had the 136V allele. In the Gibber Italicus pool, 81% of mapped reads had the 136V allele. Gibber Italicus birds are bred primarily for body shape and come in multiple color morphs, so we expect that this pool contained a mix of birds with melanin-based and melanin-loss phenotypes. Since white and lipochrome plumage in domestic canaries are recessive, it is possible that some individuals with a wild-type phenotype in a domestic population were heterozygous carriers of the 136V allele.

Interestingly, a prior study using these same pooled sequencing samples identified a variety of candidate genes associated with plumage pigment phenotypes (Bovo et al. 2023). This study did not pinpoint *EDNRB2* as a candidate gene, but did identify a significantly differentiated locus on scaffold NW_022040208.1 in several genome scans comparing melanin-loss phenotypes to melanin-based phenotypes, including Lipochrome red vs. WT, Urucum vs. WT, and White recessive vs. WT (Bovo et al. 2023). The authors identified *OCRL* as a candidate gene within this peak of genomic sequence differentiation and postulated that it may be linked to non-pigment feather traits in domestic populations. We find that *EDNRB2* (gene symbol *LOC103816487* in the canary reference genome) is within the same 700-kb haplotype as *OCRL* (see “Genes” track in fig. 4D). We hypothesize that this peak is associated with melanin loss, rather than other non-pigment traits under selection, and that *EDNRB2* is a prime candidate gene for pigment loss in Lipochrome red, Urucum, and White recessive birds.

Another prior study nominated *SCARB1* – a gene encoding a lipoprotein receptor that is important in lipid metabolism – as the gene underlying the White recessive phenotype, but these analyses compared Lipochrome yellow to White recessive, and both are melanin loss phenotypes (Toomey et al. 2017). We postulate that the purely white plumage of White recessive canaries requires mutant alleles at two loci: the D136V mutation in *EDNRB2* leads to loss of melanin pigments, and the previously characterized *SCARB1* allele is likely involved in the loss of yellow carotenoid pigment. In summary, we find evidence that *EDNRB2* may be involved in melanin loss in canaries, both in phenotypes with predominantly carotenoid plumage pigments and those with no plumage pigments at all.

### Screening reference genomes recovers two known pigmentation-associated *EDNRB2* variants in domestic species

Our alignments of *EDNRB2* orthologs identified a nonsynonymous change impacting a ligand binding residue in mallard duck *(Anas platyrhynchos)* and an in-frame insertion in swan goose *(Anser cygnoides)*, two domestic species in which prior studies have found associations between *EDNRB2* and plumage melanin loss (Li et al. 2015; Xi et al. 2020; Xi et al. 2021; Yang et al. 2022). The R332H substitution in the mallard is in transmembrane domain 6 and affects a residue predicted to play a role in ligand binding. This mutation was previously found to be associated with “spot” plumage, a partial melanin-loss phenotype (Li et al. 2015). The same R332H substitution occurs in domestic chickens, where it is associated with the “mottled” plumage phenotype, and in Japanese quail, where it is associated with “panda” plumage (Miwa et al. 2007; Kinoshita et al. 2014). Thus, this residue appears to be a mutational hotspot associated with plumage melanin loss.

The mallard annotation we used for alignment [NM_001310820.1] is derived from an individual with the previously characterized “spot” allele. Most mallard genome assemblies are from domestic Pekin ducks, which do not have the “spot” phenotype, but instead have completely white plumage. The white plumage in Pekin ducks was previously associated with an intronic insertion in *MITF* (Zhou et al. 2018). Nevertheless, we found the R332H mutation in 4 out of 5 references assembled from white-feathered Pekin ducks, as well as one assembly described as “Laying duck” and one with no listed breed (supplementary fig. S3). We would expect the *MITF* insertion allele to be epistatic to the *EDNRB2* “spot” allele; however, our identification of the *EDNRB2* R332H variant in these populations suggests that Pekin ducks may commonly harbor multiple consequential mutations that alter plumage pigmentation. Notably, the R332H mutation was not present in either genome assembly generated from wild mallards, nor was it present in two other duck species, indicating that this substitution is only found in samples with plumage melanin loss.

In the swan goose genome, we identified a three basepair insertion in the region encoding transmembrane domain 3, adjacent to a nonsynonymous substitution. The swan goose reference assembly is from a Sichuan white goose, which has no plumage melanin. Prior analysis of domestic white swan goose populations identified a 14-bp insertion in *EDNRB2* near an intron/exon boundary predicted to cause a frameshift mutation and premature stop codon (Xi et al. 2020). We found that this “new” insertion corresponds to the 14-bp insertion in exon 3 of *EDNRB2* that was identified previously. While prior work characterized the mutation as causing a frameshift (Xi et al. 2020), the NCBI annotation [XM_013199059.2] instead uses a non-canonical splice site that results in a 3-bp extension of exon 3 and the replacement of an isoleucine in the EDNRB2 consensus with a threonine and a glycine. Available reference genomes support the association of this insertion with white plumage: we find that this insertion is absent in an alternate *A. cygnoides* reference assembly from the Lion-head goose [GCA_025388735.1], which does not have white plumage. However, the insertion is present in assemblies from the white Tianfu, Zhedong, and Huoyan breeds (supplementary fig. S3), consistent with prior work showing a shared origin for white plumage across multiple breeds of Chinese domestic geese (Xi et al. 2020). Together, the swan goose and mallard mutations highlight that consequential coding mutations in *EDNRB2* can persist in avian populations without apparent negative pleiotropic effects and provide examples of how phenotype-associated mutations can be identified in reference assemblies.

In addition to the mutations in canaries, ducks, and geese discussed above, our 85-species alignment identified mutations at conserved or ligand binding residues of EDNRB2 in the Okarito kiwi (*Apteryx rowii),* budgerigar (*Melopsittacus undulatus*), ringneck pheasant (*Phasianus colchicus)* and greater honeyguide (*Indicator indicator*) genomes (table 1), as well as a single amino acid insertion in the woodland kingfisher (*Halcyon senegalensis*) genome. We believe that the coding substitution in the Okarito kiwi is unlikely to impact plumage color, as this species does not show any regional depigmentation. Understanding the potential impacts of the remaining four variants is limited by scant genetic and phenotypic data. For example, available sequencing data shows that the R188K mutation in budgerigars and the A299T mutation in ringneck pheasants are not fixed in these species (supplementary fig. S4, table S3, table S4). Prior work has documented variable pigmentation, including melanin loss phenotypes in both budgerigars and ringneck pheasants (Bruckner 1941; Martin 2002; Cooke et al. 2017; Kayvanfar et al. 2017); however, it is impossible to tell if the mutations identified here are associated with loss of plumage pigment as phenotype information is not available for sequenced samples. While the greater honeyguide has white ventral plumage that could be driven by an *EDNRB2* mutation, it is not possible to infer from a single sequencing sample if this mutation is associated with plumage pigmentation more broadly. In the woodland kingfisher genome [GCA_013397595.1], the amino acid insertion is caused by an intronic single nucleotide polymorphism that creates an additional splice acceptor (supplementary fig. S4), but it is not clear if this splice acceptor is actually used. Future projects using larger samples with known phenotypes from these additional species, or sequencing closely related species with varying phenotypes, could shed light on the impact of these candidate *EDNRB2* variants on plumage pigmentation.

### *EDNRB2* coding variation may explain pigmentation phenotypes in cockatoos

To complement the strategy provided by our 85-species gene alignments, we next pursued targeted analyses of a smaller group of closely related species. We looked for lineages where whole-genome sequencing was available for multiple species that share similar melanin loss phenotypes and other close relatives that do not show melanin loss. With these criteria in mind, we chose to screen genome assemblies from cockatoos (family Cacatuidae) for *EDNRB2* variation.

Both wild and domestic cockatoo populations show melanin loss. Notably, all species in the genus *Cacatua* show total loss of plumage melanin, though some have regional red or yellow psittacofulvin (a parrot-specific pigment) coloration (fig. 5A). Thirteen cockatoo genomes have been sequenced: nine *Cacatua* species with no plumage melanin, 3 species with regional melanin loss, and 1 species with no loss of plumage melanin. Most cockatoo genomes are not annotated. We used a user-submitted genome annotation to identify full-length *EDNRB2* coding sequences for 10 out of 13 sequenced cockatoo species; the remaining three genomes were highly fragmented and full length *EDNRB2* coding sequences could not be reconstructed. We aligned the 10 full-length cockatoo *EDNRB2* coding sequences and found an increased dN/dS ratio in several branches within the *Cacatua* clade (fig. 5A). We identified 23 nonsynonymous substitutions throughout the coding sequence and one deletion in the region encoding the extracellular N-terminal protein domain. Three of the 23 nonsynonymous substitutions mapped to amino acid positions that that were fully conserved in our tetrapod endothelin receptor alignment (fig. 5B); all 3 of these amino acid substitutions were predicted to be damaging by PolyPhen2. One of these substitutions, A320V in transmembrane domain 6, is shared by the seven *Cacatua* species, all of which have lost all plumage melanin. The other two mutations at conserved sites, C118S in the intracellular domain between transmembrane domains 2 and 3 and T175A in transmembrane domain 3, are specific to the galah, which shows regional loss of plumage melanin. The cockatiel, which has regional melanin loss, and palm cockatoo, which has no regional melanin loss, do not have any substitutions at fully conserved or ligand binding residues.

**Figure 5:**
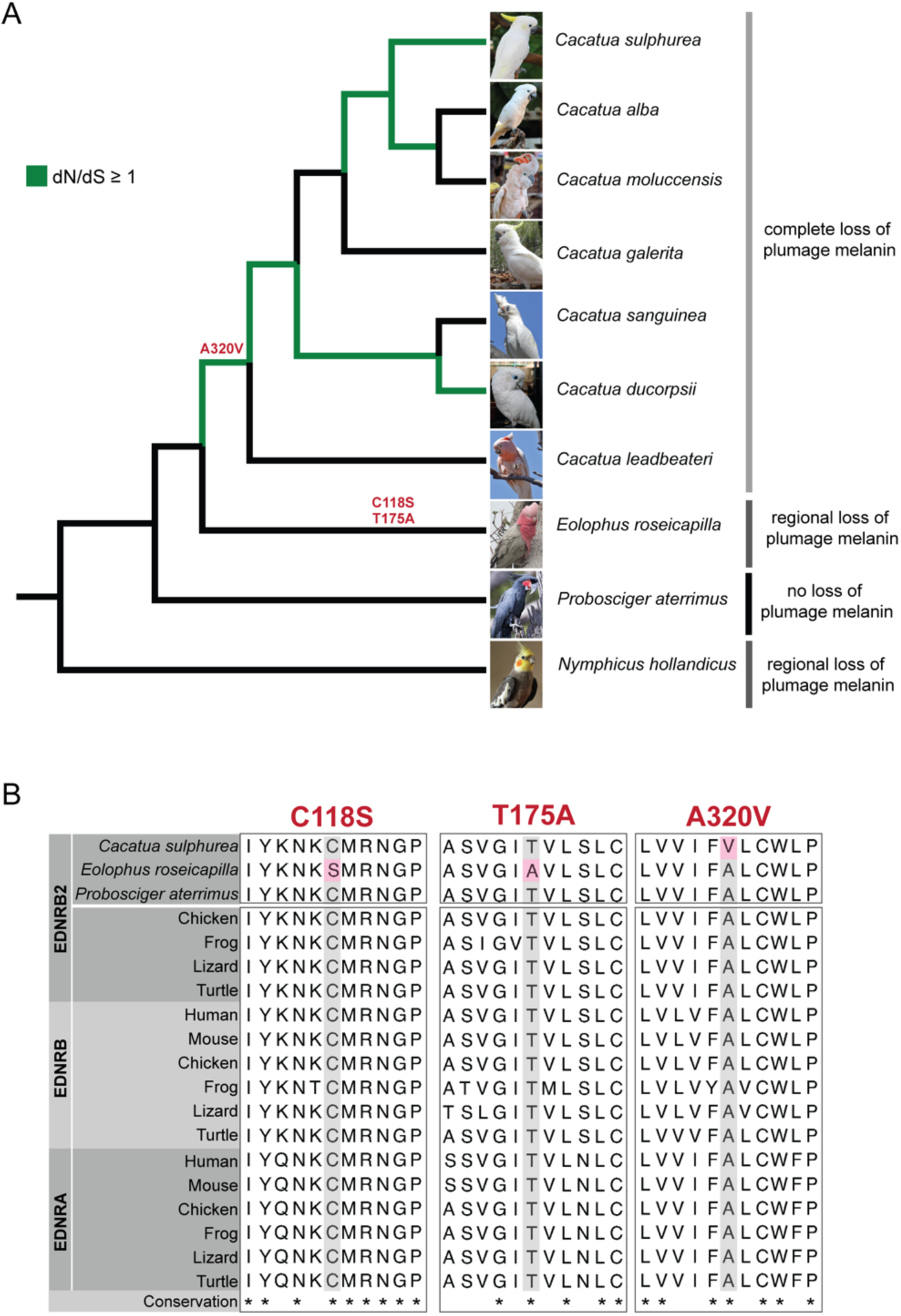
Comparative analysis of cockatoo *EDNRB2* coding sequences identifies multiple candidate substitutions that may drive melanin loss. (A) Cladogram illustrating the relationships between 10 cockatoo species. Branches with a dN/dS ratio greater than 1 are highlighted in green. Images illustrate pigment phenotypes for each species, image credits and creative commons license details are in supplementary table S6. The substitutions identified at three highly conserved residues are noted in red above the branch where they occurred. (B) Regional alignments highlighting the nonsynonymous substitutions identified in cockatoo genomes (pink) compared to the reference genomes of additional cockatoo species and a multi-species alignment of EDNRA, EDNRB, and EDNRB2 from diverse vertebrates. * indicates amino acids that are 100% conserved in the vertebrate endothelin receptor alignment.

While we were not able to reconstruct the full *EDNRB2* coding sequence for the gang-gang cockatoo (*Callocephalon fimbriatum*), we confirmed that this species does not carry any of the above cockatoo mutations at conserved residues. This is consistent with phenotype, as the gang-gang cockatoo is sexually dimorphic and shows melanin loss only on the heads of males. Both the *Cacatua* species and galah have more extensive and non-dimorphic melanin loss. We hypothesize that the A320V mutation may contribute to global loss of plumage melanin in *Cacatua* species, while the C118S and T175A mutations may underlie regional melanin loss in the galah. While this candidate gene approach has not ruled out contributions to these phenotypes from other pigment genes, our findings suggest that damaging *EDNRB2* mutations are associated with plumage pigment variation in some wild species.

## DISCUSSION

We examined *EDNRB2* coding changes across a diverse array of avian species. We found that, compared to *EDNRB, EDNRB2* has a significant increase in the number of total nucleotide substitutions and nonsynonymous substitutions, indicating differences in the selective pressures that act on each of these genes. The specialized functions of these two endothelin B receptor genes likely underlie some of the differences in observed mutational burden. *EDNRB* is critical for migration of skeletal and trunk neural crest cells, and loss of function mutations are likely to be incompatible with reproductive success or survival in the wild, while *EDNRB2* mutations are expected to cause changes in plumage color but not incur major pleiotropic effects impacting viability (Pla and Larue 2003; Square et al. 2016; Liu et al. 2019; Square et al. 2020).

Through our analyses of *EDNRB2* coding sequences in avian genome assemblies, we identified both novel and previously characterized coding variants in domestic species that are associated with melanin loss, and coding variants in wild species that may underlie changes in plumage melanin (fig. 6). Here, we focused on variation at highly conserved amino acid residues involved in ligand binding. Many previously identified mutations also affect conserved or ligand binding residues (fig. 6). However, prior work in domestic species has also identified structural variation, noncoding variants, and coding variation at less conserved sites in *EDNRB2* that are associated with melanin loss (Miwa et al. 2007; Kinoshita et al. 2014; Li et al. 2015; Xi et al. 2020; Xi et al. 2021; Rodríguez-Martínez 2022; Maclary et al. 2023). These other types of variation are more difficult to evaluate using existing reference genomes and sequencing data. In particular, noncoding regions are less conserved among species and many published avian genomes are highly fragmented, making alignment of noncoding sequence and identification of possible structural variants especially challenging.

**Figure 6.**
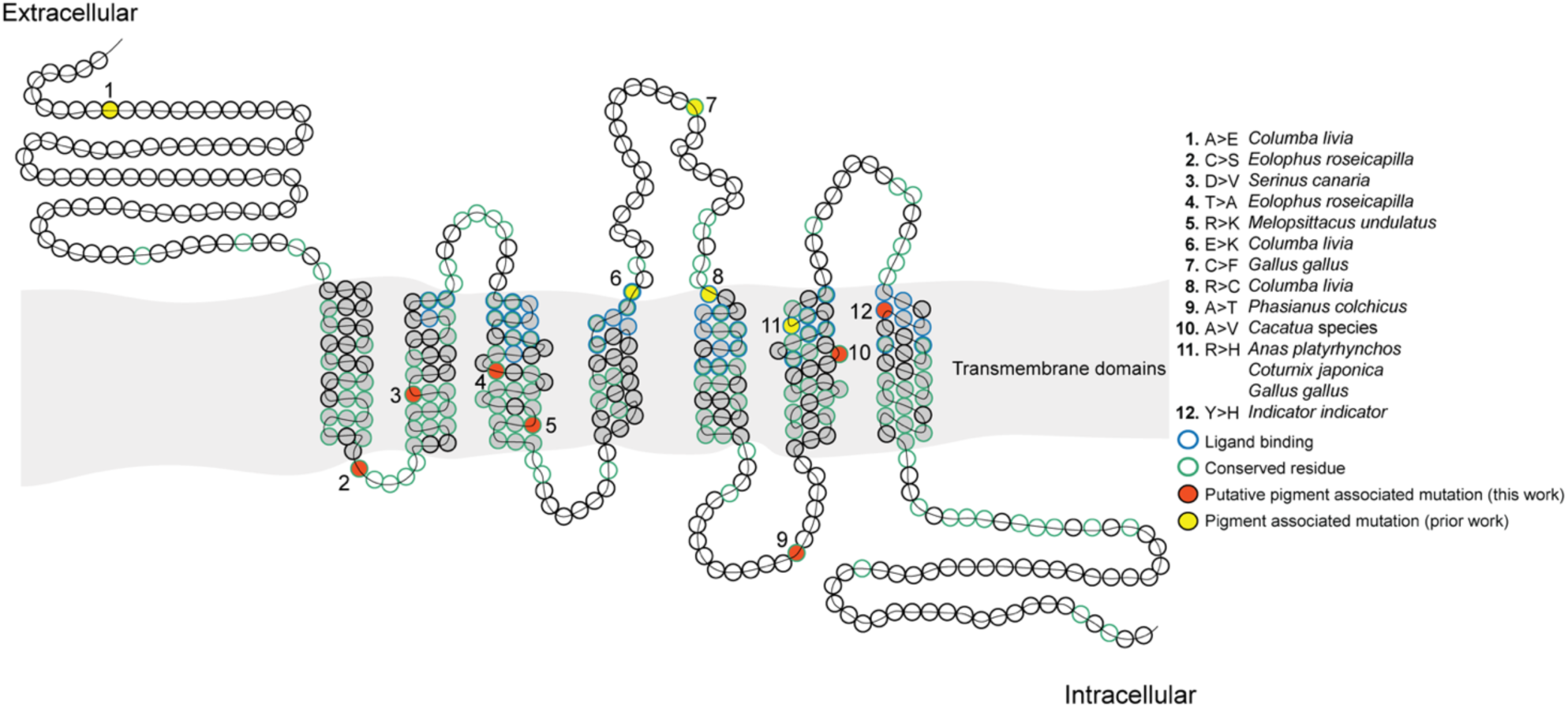
Diagram illustrating the locations of EDNRB2 variants associated with plumage pigmentation. Each circle represents an amino acid, starting with position 1 on the left and ending with position 436 on the right. A continuous black line shows the linear order of amino acids throughout the structure diagram. Transmembrane domains are indicated in gray, amino acids predicted to be involved in ligand binding are outlined in blue. Amino acids that are 100% conserved in our multi-species alignment of EDNRA, EDNRB, and EDNRB2 are outlined in green. Sites with putative pigment associated variants identified in this work are shown in orange. Previously published pigment-associated coding variants are indicated in yellow. Sites with pigment-associated variants are numbered for identification.

Structural and noncoding variation may be common at the *EDNRB2* locus due to the nature of the DNA sequence in the region. We previously found that repeats and transposable elements were enriched at the *EDNRB2* locus in domestic pigeon (Maclary et al. 2023), and *EDNRB2* resides on a chromosome that has undergone recurrent major structural rearrangements in parrots (Huang et al. 2022). Some of these rearrangements may have directly affected the *EDNRB2* locus. For example, we found that the budgerigar genome has multiple pseudogenized endothelin B receptors within a 28-Mb window on chromosome 6 that overlaps the *EDNRB2* locus (Gene IDs LOC117436282, LOC117436268, LOC117436259, LOC117436264, and LOC117436273). Targeted sequencing of specific clades with polymorphic melanin phenotypes could reveal the prevalence of noncoding and structural variation at the *EDNRB2* locus and, as available genome assemblies expand in number and improve in quality, future work can better evaluate possible structural or noncoding variation across species. In this study, we identified candidate coding variants that may drive plumage variation in a small number of species, and we hypothesize that a mix of coding and regulatory variation at *EDNRB2* may underlie plumage pattern changes in far more species.

### Pleiotropy may constrain the genetic mechanisms that drive plumage pigment diversity

We compared synonymous and nonsynonymous substitution rates between *EDNRB2* and five other genes commonly associated with polymorphic melanin-based coloration in domestic avian species. We found variation in nonsynonymous and synonymous mutation rate, and this variation supported the hypothesis that pleiotropy may constrain the genes and types of mutations that drive variation in plumage pigment in birds. We found that two genes with roles in both pigmentation and crucial events in embryonic development, *MITF* and *SOX10*, showed significantly reduced numbers of nonsynonymous and total substitutions per 100 codons. Three genes with fewer known developmental pleiotropies – *EDNRB2, MC1R,* and *TYRP1* – had significantly higher numbers of both total and nonsynonymous substitutions when normalized for length, and *ASIP* had both increased dN/dS values per branch and significantly higher numbers of mutations per 100 codons compared to all other pigment genes assayed.

Several studies, including both candidate gene approaches and unbiased genome-wide screens, have implicated *MC1R* in plumage pigment variation in both domestic and wild avian species (Mundy 2005; Guernsey et al. 2013; Corso et al. 2016; Campagna et al. 2022; Ono et al. 2024). As *EDNRB2* shows comparable substitution rates and lacks clear pleiotropic effects, we anticipate that *EDNRB2* is similarly likely to underlie variation in plumage melanin across a wide array of species. Prior work has also shown that many types of changes at the *EDNRB2* locus underlie melanin-loss phenotypes without known deleterious pleiotropic effects, including coding changes, regulatory changes, structural variants, and even complete deletion of the *EDNRB2* gene (Xi et al. 2020; Rodríguez-Martínez 2022; Maclary et al. 2023). As a result, we expect few constraints on the types of *EDNRB2* mutations that could contribute to changes in plumage melanin across avian species.

### Regional loss of melanin pigment plays a critical role in establishing non-melanin pigment patterns

*EDNRB2* mutations in domestic species are typically thought to be associated with white plumage. We identified mutations in two species, canaries and cockatoos, where feathers that lack melanin still have other non-melanin pigments. Non-melanin pigments in birds include carotenoids and psittacofulvins, which are responsible for bright yellow, orange, red, and pink hues (Price-Waldman and Stoddard 2021). Psittacofulvins are found only in parrots and are endogenously synthesized rather than obtained through diet. Prior work has found that melanin-based and non-melanin colors can combine to produce colors and hues that neither type of pigment can produce alone, but it is unclear whether there is coordinated control of melanin and non-melanin pigments, nor do we understand completely how the genetic and biochemical pathways governing melanin-based and non-melanin colors interact to create both a wide array of colors and complex patterns.

One recent comparative genomic study of yellow-shafted and red-shafted flickers identified associations between carotenoid traits and melanogenesis genes (Aguillon et al. 2021). Three possible explanations were proposed for this novel finding: first, melanin is present in the carotenoid-containing feathers that define the trait but it is masked by carotenoids; second, the genes involved in melanin pigmentation are pleiotropic and also play roles in production or deposition of carotenoid pigmentation that have yet to be characterized; or third, downregulation of melanin genes is required to prevent masking of carotenoid pigments. We find this third explanation, that disruption of melanin pigmentation is required for the visibility of some carotenoids, to be the most compelling. Our findings in canaries and cockatoos suggest that *EDNRB2*-mediated melanin loss could be critical for establishing non-melanin based plumage patterns in both wild and domestic species.

In canaries, the Lipochrome yellow and Lipochrome red colors in domestic breeds rely on loss of melanin to create a bright, uniform plumage color. Our analysis finds that these birds carry the same *EDNRB2* mutation as the completely depigmented White recessive canary. Loss of melanin differentiates the domestic lipochrome colors from the duller yellow-green and brown plumage color of wild common canaries, whose feathers contain melanin (Lopes and Carneiro 2024). While few wild songbirds exhibit total melanin loss like domestic Lipochrome canaries, many do have regional patches of brilliant yellow, red, or orange plumage that is reminiscent of these lipochrome colors. We hypothesize that changes in melanin production and distribution are essential for establishing many of the carotenoid-based plumage colors and patterns seen in songbirds and other clades. For example, prior work examining the pigment types used in the orange, yellow, and red feathers of orioles found that some species produce these colors solely from carotenoids, some solely from pheomelanin, and others a mix of both carotenoids and melanins (Hoffman et al. 2007). Therefore, these species use a mix of carotenoids and melanin pigments to titrate the hue and intensity of feather colors.

The interactions between melanin and non-melanin pigments are not limited to carotenoid traits. We identified putative damaging *EDNRB2* mutations in cockatoos, which have bright regions of red, yellow, or pink feathers produced by psittacofulvin pigments. Two previous studies of domestic Psittaciformes provide insight into the interaction between melanins and psittacofulvins. Prior work on budgerigars identified a polyketide synthase, MuPKS, that is required for production of yellow pigment (Cooke et al. 2017). Wild-type budgerigars have green and yellow plumage, and loss-of-function mutations in MuPKS produce birds that have blue and white plumage (Cooke et al. 2017). The blue coloration in feathers is structural, resulting from dispersion of light by structures within the feather, including both keratins and melanin granules. Thus, some structural colors are melanin-dependent (Price-Waldman and Stoddard 2021; Saranathan and Finet 2021). In parrots, green feathers contain melanin and arise from a mix of blue structural color and yellow psittacofulvin, while yellow patches contain only psittacofulvin. Yellow feathers become white and green feathers become blue when psittacofulvin synthesis is inhibited (Cooke et al. 2017; Neves et al. 2020; Price-Waldman and Stoddard 2021).

Another recent study examining a sex-linked “yellow” phenotype in three species of domestic *Psittacula* parakeets similarly supports the role of melanin loss in mediating complex plumage patterns. *SLC45A2* has been linked to melanin reduction or loss in several species, including humans, horses, chickens, quail, and pigeons (Cieslak et al. 2011; Reissmann and Ludwig 2013; Domyan et al. 2014; Roy et al. 2023). In parakeets, wild-type individuals have regions of green, blue, purple, or black feathers. *SLC45A2* loss-of-function mutants instead have only yellow or red feathers (Roy et al. 2023). These studies show that melanin pigments, melanin-based structural colors, and psittacofulvins in different combinations and quantities work together to produce the rainbow palette that characterizes parrot species. While melanin production is crucial for some plumage colors, regional melanin loss is equally important for establishing the bright plumage with solely psittacofulvin-based coloration, such as the yellow plumage patches in wild budgerigars.

Melanin-free plumage patches can be evolutionarily important, including bright patches of carotenoid coloration that are thought to serve as “honest signals” of fitness in many species (Hill 1991). We conclude that regional melanin loss is critical for multiple aspects of plumage coloration across avian species, not only the development of white plumage. Our analysis suggests that variation at the *EDNRB2* locus could play a key role in both establishing white plumage patches and in setting the stage for the development of bright non-melanin plumage colors across avian species.

## METHODS

### Identification of gene orthologs in annotated genomes

*EDNRB2*, *EDNRB1, ASIP, MC1R, MITF, SOX10,* and *TYRP1* orthologs in NCBI-annotated genomes were identified using a combination of the NCBI ortholog database and nucleotide blast of NCBI nr/nt sequence databases. We extracted the coding portion of mRNA sequences from NCBI accessions (supplementary table S1) using NCBI batch entrez, selecting “Coding Sequences” in FASTA Nucleotide format.

For *EDNRB2* and *EDNRB1* sequences from submitter annotated genomes, we used the provided GFF files to extract coding sequences from genomic FASTA files using gffread. We found that some annotations were truncated by 1-3 amino acids, in which case we extended the extracted sequences to the nearest in-frame methionine. Annotations that were manually extended are noted in supplementary table S1.

Several orthologs of *MC1R* – a single-exon gene – in avian genomes are annotated by NCBI as fusions with the adjacent *TUBB3* gene. In these cases, the first exon corresponds to the *MC1R* coding sequence and subsequent exons are part of the *TUBB3* gene. These fusion annotations are spliced at the normal “stop” codon for the *MC1R* sequence, and are prevalent in genome databases due to intergenic splicing that produces chimeric *MC1R-TUBB3* transcripts detectable in transcriptome datasets (Herraiz et al. 2017). For annotations where sequence for *MC1R* and *TUBB3* are fused, we used the sequence encoding the first 314 amino acids for *MC1R* alignments. *MC1R* sequences derived from *TUBB3* annotations are indicated in supplementary table S1.

### Coding sequence alignment and variant identification

The coding portion of mRNA sequences were downloaded from NCBI by accession using Batch Entrez. Sequences were aligned with PRANK (v.170427), using the “-codon” option for codon-aware alignment (Löytynoja 2013). Sequence alignments were visualized using Jalview (Waterhouse et al. 2009).

Cladograms for the 85 species and 35 species alignments were constructed based on published molecular phylogenies. Family-level trees were based on the consensus tree from Braun and Kimball’s re-analysis of multiple data types (Braun and Kimball 2021). Within-family relationships were reconstructed from multiple sources (Brown and Toft 1999; Sun et al. 2017; Oliveros et al. 2019; Kuhl et al. 2020; Chen et al. 2021; Smith et al. 2022). PRANK alignments were analyzed in HyPhy v. 2.5.58 (Pond et al. 2019)(www.hyphy.org) with these cladograms as the input tree for all genes. The “FitMG94.bf” and “AncestralSequences.bf” functions were used to identify synonymous substitutions, nonsynonymous substitutions, and local dN/dS for each cladogram branch (https://github.com/veg/hyphy-analyses/). To normalize numbers of synonymous and nonsynonymous substitutions across genes with varying lengths for comparison, we calculated the number of synonymous and nonsynonymous substitutions per 100 codons. We compared substitution counts and dN/dS values between genes using two-tailed t-tests.

Translated coding sequences for EDNRB and EDNRB2 ranged in length, with extensive sequence variation in the extracellular N-terminus of the protein prior to the start of the first transmembrane domain. For alignments, all coordinates were standardized to the *Gallus gallus* reference, since chicken is a commonly used model with a well-curated annotation set. Three out of 85 EDNRB annotations and 39/85 EDNRB2 annotations had N-terminal extensions beyond the chicken annotation that were excluded from analysis. In addition, 15/85 EDNRB annotations were truncated compared to chicken. Truncated annotations start before the first transmembrane domain but a shorter annotated extracellular N terminus. Consequently, consensus sequences and substitutions prior to chicken M87 in EDNRB were identified from a reduced number of samples.

### Identification of conserved domains and residues

To identify conserved residues, we aligned protein sequences available in NCBI databases for three endothelin receptors (EDNRA, EDNRB, and EDNRB2) from human (*Homo sapiens*), mouse (*Mus musculus*), chicken *(Gallus gallus*), western clawed frog (*Xenopus tropicalis*), common wall lizard (*Podarcis muralis),* and green sea turtle (*Chelonia mydas*). Accessions are listed in supplementary table S1. Protein sequences were aligned using Clustal Omega (Madeira et al. 2019), and alignments visualized using Jalview (Waterhouse et al. 2009). Transmembrane and ligand binding domains in EDNRB and EDNRB2 were identified based on the NCBI Conserved Domain Database accessions cd15976 and cd15977 (Lu et al. 2019).

### Targeted assessment of coding variation in reference assemblies and sequencing datasets using tblastn

For mutations identified in the *Serinus canaria, Anas platyrhynchos, Anser cygnoides,* and *Phasianus colchicus* genomes, we screened additional genome assemblies (see accessions in supplementary table S4) and samples in the NCBI Sequence Read Archive (SRA) (see accessions in supplementary table S5) for the variants of interest using NCBI tblastn (Johnson et al. 2008). We used tblastn to compare EDNRB2 coding sequence to whole-genome shotgun contigs (wgs) or samples from the Sequence Read Archive (SRA) from the same species or closely related species as the search database. We then examined alignment results to assess the presence or absence of variants of interest.

### Alignment and analysis of *Serinus canaria* whole genome sequencing

Publicly available sequencing data for *S. canaria* was downloaded from the NCBI SRA using sra-toolkit (https://github.com/ncbi/sra-tools). Accessions for all previously published SRA datasets are in supplementary table S3. Fastq files were aligned to the *S. canaria* reference genome (GCA_022539315.2) using bowtie2 with the "--local” setting (Langmead and Salzberg 2012).

SNPs were called from alignments using bcftools mpileup (v. 1.16) (Danecek et al. 2021), and genotypes plotted across the candidate region using the R packages “vcfR” (Knaus and Grünwald 2017) and “GenotypePlot” (https://github.com/JimWhiting91/genotype_plot). Genotypes for the candidate SNP of interest (NC_066318.1:18,571,511) in each sample pool and allelic ratios for pools with both alleles were determined from both SNP calls and manual examination of BAM read alignments in IGV (Robinson et al. 2011).

### Alignment and analysis of *Melopsittacus undulatus* RNA-seq

Publicly available sequencing data for *M. undulatus* was downloaded from NCBI using sra-toolkit (https://github.com/ncbi/sra-tools). Accessions for previously published RNA-seq datasets are in supplementary table S5. Fastq files were aligned to the *M. undulatus* reference genome (GCF_012275295.1) using STAR (Dobin et al. 2013). Genotypes at the variant of interest (NC_047532.1:73,663,884) for each sample were determined by visualizing BAM format alignments in IGV (Robinson et al. 2011).

### Identification of mRNA sequence from unannotated *Cacatuidae* genomes

To obtain coding sequences from unannotated *Cacatuidae* genomes, we extracted DNA sequence for individual exons of *EDNRB2* (EOLROS_R08102) from the *Eolophus roseicapilla* reference genome (GCA_013397615.1) using Bedtools getfasta. The *EDNRB2* protein sequence obtained using Bedtools was partial, lacking a starting methionine. We extended the 5’ end of the EOLROS_R08102 annotation by three base pairs, which produced a coding sequence initiating with a methionine. We used NCBI the tblastn web service to align *E. roseicapilla* exons to whole-genome shotgun contigs for *Cacatua, Callocephalon, Probosciger*, and *Nymphicus* species. We identified complete coding sequences for the palm cockatoo (*Probosciger aterrimus),* cockatiel (*Nymphicus hollandicus),* and 7 *Cacatua* species (white cockatoo, *C. alba*; yellow-crested cockatoo, *C. sulphurea*; Solomons corella, *C. ducorpsii*; sulphur-crested cockatoo, *C. galerita*; pink cockatoo, *C. leadbeateri*; little corella, *C. sanguinea*; and salmon-crested cockatoo, *C. moluccensis*). Accessions for all genome assemblies used are in supplementary table S4.

## DATA AVAILABILITY

Newly-generated data underlying this article are available in the article and its supplementary material. Previously published genomes, annotations, and sequencing data analyzed in this article are publicly available in NCBI repositories. All accessions are listed in supplementary tables S1 (gene and protein accessions), S4 (genome assembly accessions), and S5 (SRA dataset accessions).

## ACKNOWLEDGEMENTS

We thank Jim Baldwin-Brown, Olivia Inman, and Atoosa Samani for technical and conceptual discussion over the course of this project and Yangsu Ren for critical reading of the manuscript. The support and resources from the Center for High Performance Computing at the University of Utah are gratefully acknowledged.

## AUTHOR CONTRIBUTIONS

E.T.M. and M.D.S. designed the study. E.T.M. performed all data analysis and visualization. M.D.S. supervised the study. E.T.M. and M.D.S. wrote the manuscript.

