## Supplementary figures 1-4 for "Widespread variation in *EDNRB2* is associated with diverse melanin loss phenotypes across avian species"

Emily T. Maclary* and Michael D. Shapiro
School of Biological Sciences, University of Utah, Salt Lake City UT USA

*Author for Correspondence
Emily T. Maclary, School of Biological Sciences, 257 South 1400 East, Salt Lake City, UT 84112 USA; phone +1 801 551 6812;

Figure S1
Figure S2
Figure S3
Figure S4

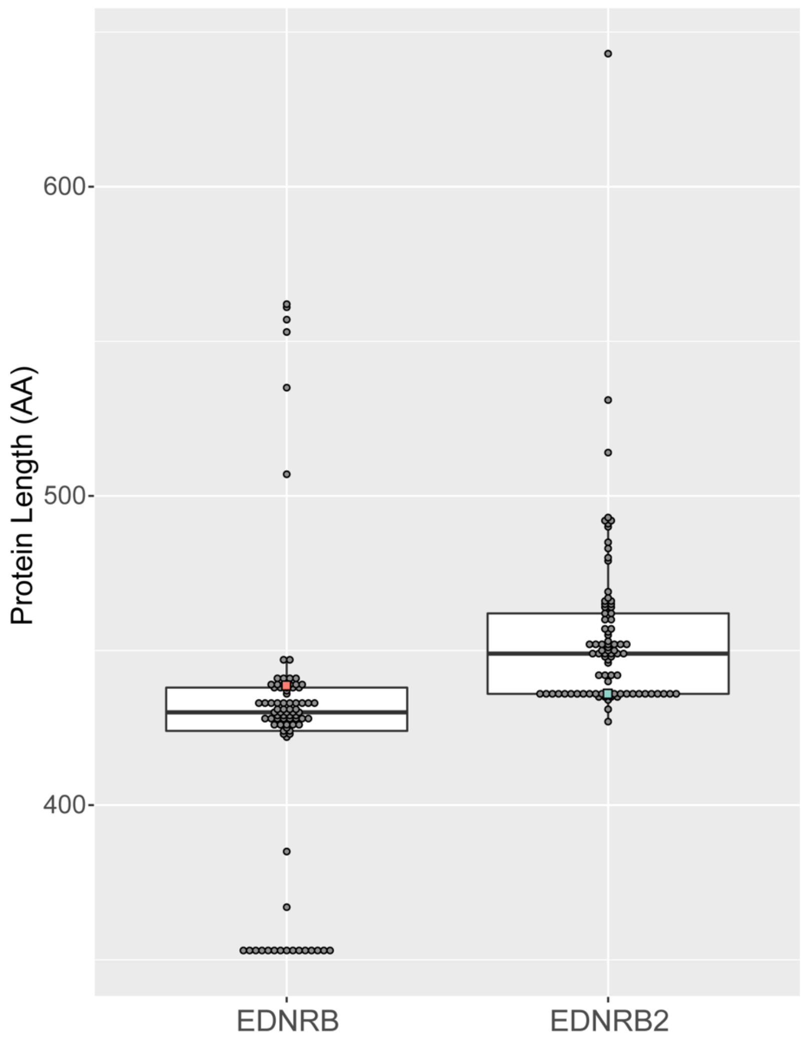


**Figure S1. Protein length varies between species in EDNRB and EDNRB2 annotations.** Overlaid boxplots and dotplots for EDNRB (left) and EDNRB2 (right) depicting the range in annotated protein length. Y axis, protein length (amino acids). Each gray dot represents a species in the 85-species alignment. Red and green squares represent the chicken EDNRB (red) and EDNRB2 (green) sequences used as a reference for subsequent analysis. Boxes span from the first to third quartile of each data set, with lines indicating the median. Whiskers span up to 1.5x the interquartile range.

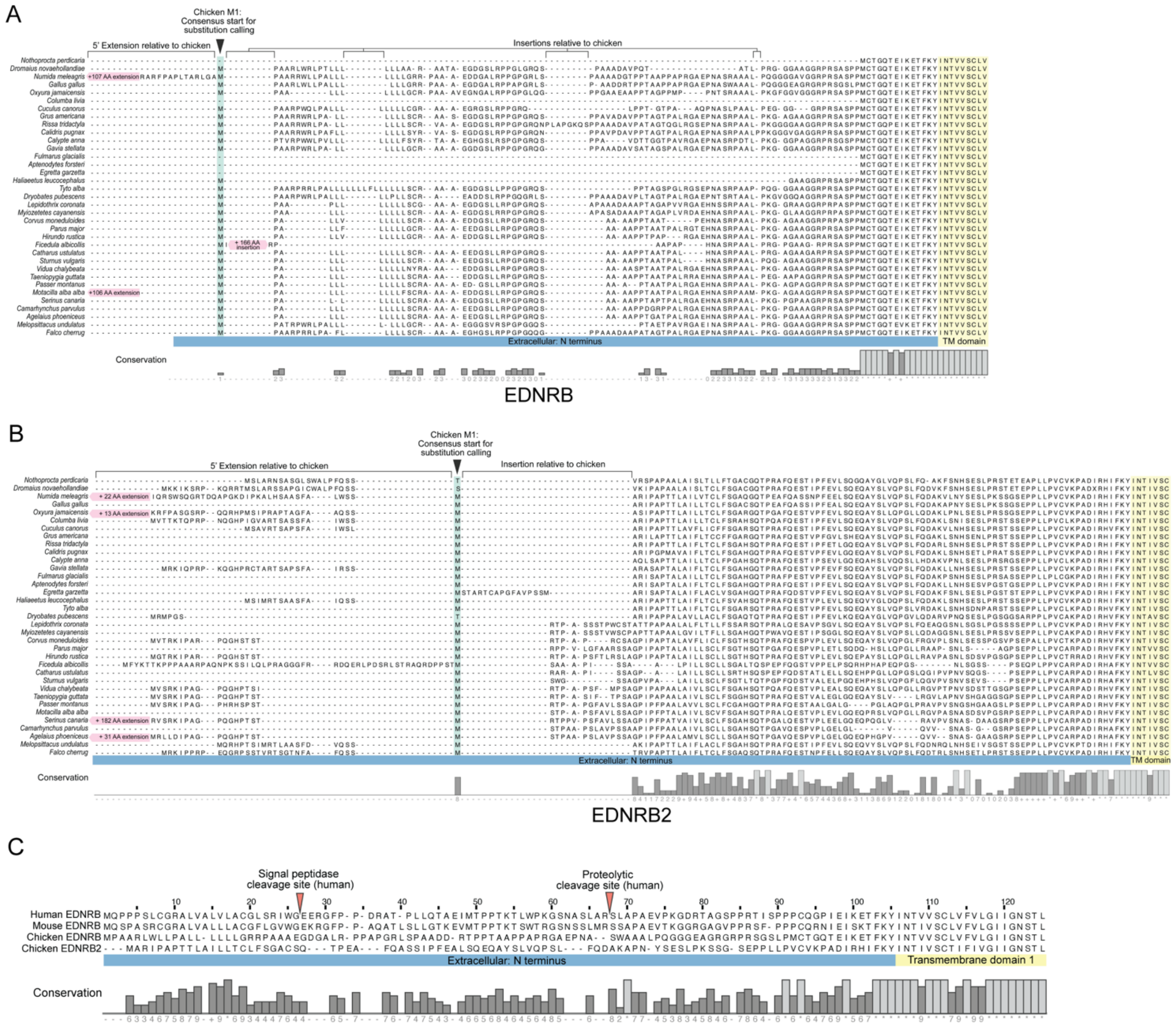


**Figure S2. The extracellular N-terminal domains of EDNRB and EDNRB2 are highly variable between species.** (A-B) 35-species alignment illustrating N-terminal variation in EDNRB (A) and EDNRB2 (B). Along the X axis, colors mark the boundaries of the extracellular portion of the protein (blue) and the start of the first transmembrane domain (yellow). (C) Alignments of the N-terminus of human, mouse, and chicken EDNRB and Chicken EDNRB2. Red arrows mark the known signal peptidase cleavage site and proteolytic cleavage site in humans. Along the X axis, colors mark the boundaries of the extracellular portion of the protein (blue) and the start of the first transmembrane domain (yellow).

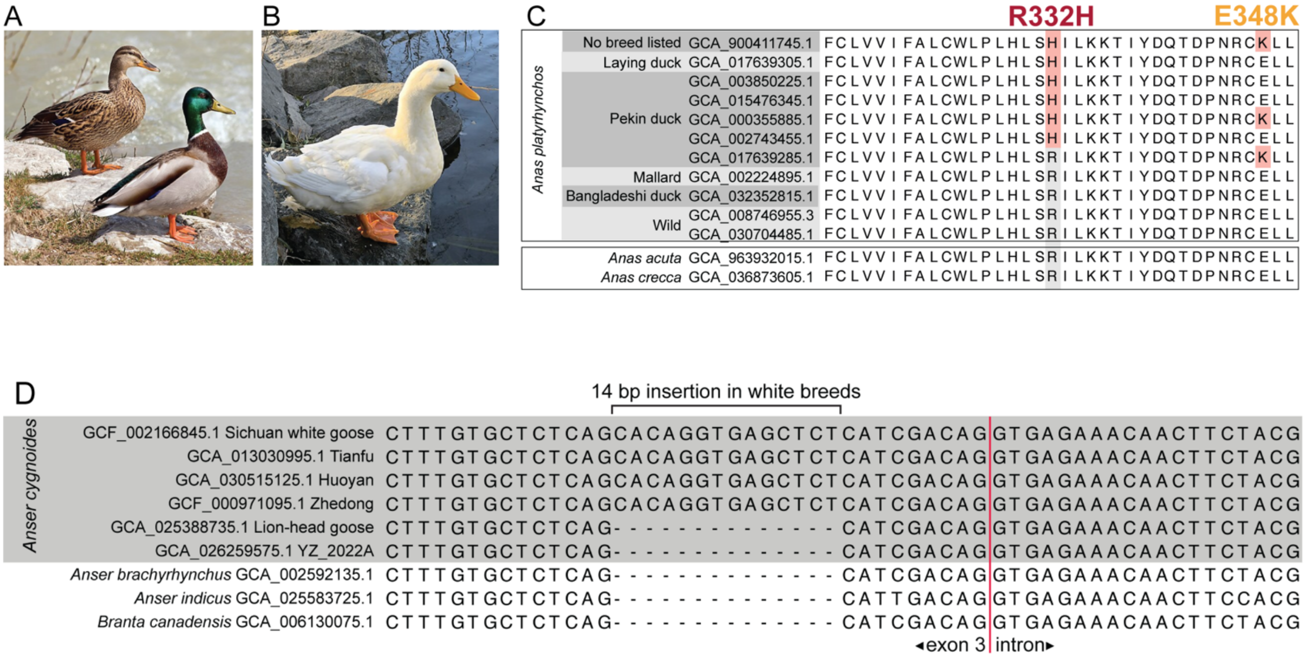


**Figure S3. Evaluation of EDNRB2 variants in *Anas platyrhynchos* and *Anser cygnoides*.** (A) Wild-type *Anas platyrhynchos* are sexually dimorphic and have melanin-based plumage colors. (B) A representative Pekin duck (domestic form of *A. platyrhynchos*). Image credits and creative commons license details are in table S6. (C) Alignments of amino acid sequence from 11 *A. platyrhynchos* genomes and two additional *Anas* species for the region containing the R332H substitution. The H variant and a second nonsynonymous variant present in a subset of *A. platyrhynchos* genomes, E348K, are highlighted in red. (D) Nucleotide alignments from six *Anser cygnoides* genomes and three additional goose species showing a 14 basepair insertion in four *A. cygnoides* genomes. A red line marks the intron-exon boundary at the end of exon 3.

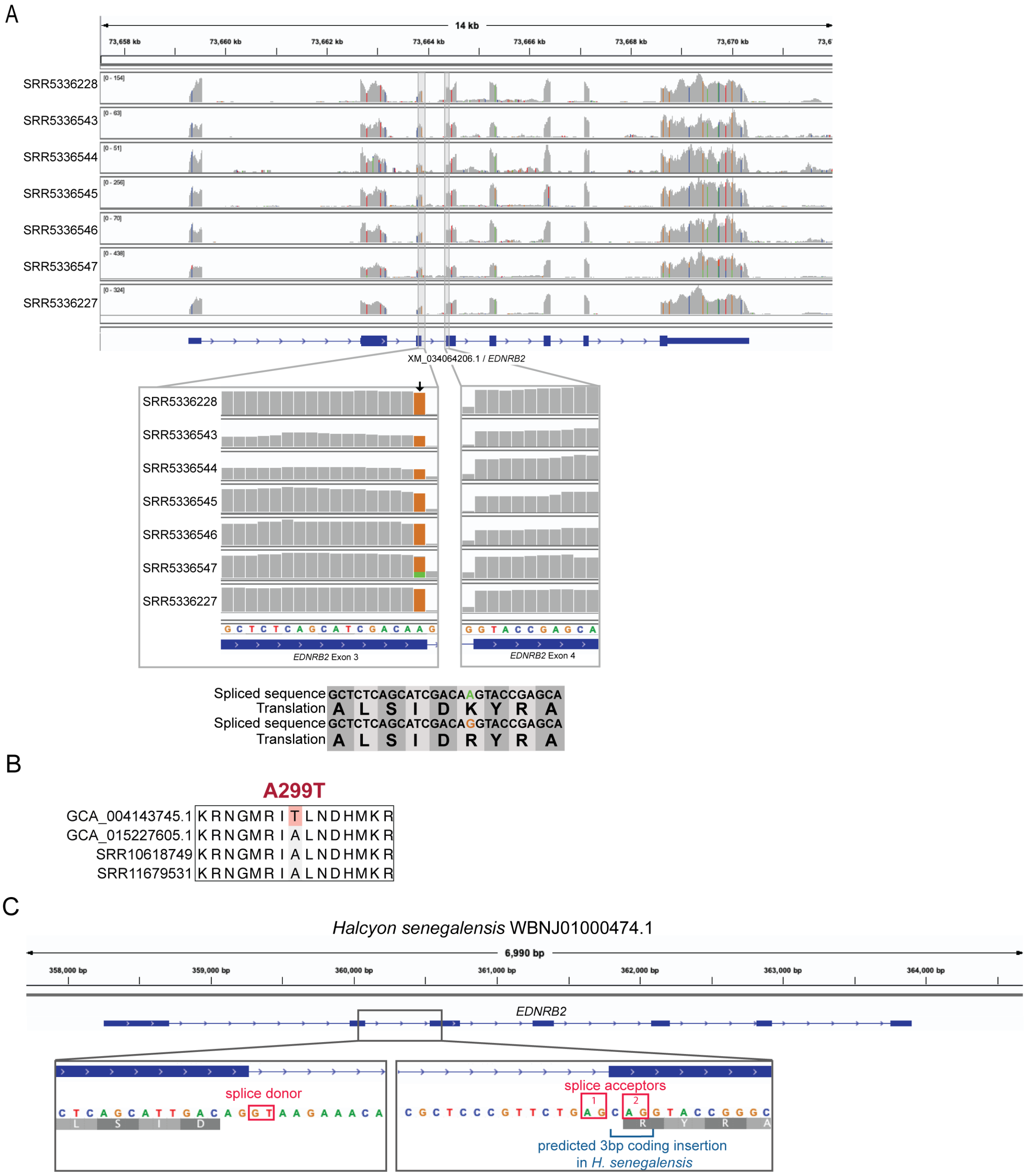


**Figure S4. Characterization of *EDNRB2* variants identified in pheasant, budgerigar, and kingfisher genomes** (A) Coverage tracks illustrating RNA-seq alignments from six *Melopsittacus undulatus* samples at the *EDNRB2* locus. The Y axis of each sample track shows sequencing depth, and the X-axis genome location. Colored lines indicate sites where the nucleotide present in aligned reads differs from the reference genome. A diagram of the *EDNRB2* gene is shown below. Inset images below zoom in on exons 3 and 4. An arrow marks a single nucleotide polymorphism at the boundary of exon 3. The reference genome has an "A"

at this site. Six out of seven RNA-seq samples have a “G” at this site, indicated by the orange bar. The seventh is heterozygous, indicated by the split orange and green bar. Spliced sequence and translations are shown below. The “A” allele produces a lysine (K) at position 188 in EDNRB2, while the “G” allele produces an arginine (R). (B) Alignments of amino acid sequence from *Phasianus colchicus* samples for the region containing the A299T substitution. The T variant, highlighted in red, is present in the current reference genome (GCA_004143745.1), but not in three other samples assessed. (C) *EDNRB2* gene model and nucleotide sequence from the *Halcyon senegalensis* reference genome. Zoomed regions show the exon 2/intron 2 boundary (left) and the intron 2/exon 3 boundary (right). Red boxes highlight possible splice donor and splice acceptor sites. There are two possible splice acceptors in this region in *H. senegalensis*; the annotation uses splice acceptor 1, which produces a 3bp coding insertion in the mRNA sequence (blue bracket) compared to other avian *EDNRB2* coding sequences.
